# Prediction of Carbon Emissions in Guizhou Province-Based on Different Neural Network Models

**DOI:** 10.1101/2023.12.21.572929

**Authors:** Lian Da, Yang ShiQiang, Yang Wu, Ran WenRui, Zhang Min

## Abstract

Global warming caused by greenhouse gas emissions has become a major challenge facing people all over the world. The study of regional human activities and their impacts on carbon emissions is of great significance to achieve the ambitious goal of carbon neutrality and sustainable economic development. Guizhou Province is a typical karst area in China, and its energy consumption is mainly based on fossil fuels.Therefore, it is necessary to predict and analyze its carbon emissions. In this paper, BP neural network and extreme learning machine (ELM) model, which have the advantage of nonlinear processing, will be used to predict the carbon emissions of Guizhou Province from 2020 to 2040. Based on the energy consumption data of Guizhou Province, the carbon emissions of Guizhou Province are calculated by using the conversion method and the inventory compilation method. The data show that the carbon emissions of Guizhou Province show an “S” growth trend; In this paper, 12 influencing factors of carbon emissions are selected, and five influencing factors with larger correlation are screened out by using grey correlation analysis method, and the prediction model of carbon emissions in Guizhou Province is established and simulated, and the prediction performance of BP neural network, ELM and WOA-ELM are compared respectively. Compared with ELM model and BP neural network model, the prediction accuracy of WOA-ELM model is higher; Finally, three development scenarios of carbon emissions are set by scenario analysis, which are baseline scenario, high-speed scenario and low-carbon scenario. On this basis, the size and time of peak carbon emissions in Liaoning Province from 2020 to 2040 are predicted based on WOA-ELM model. The results show that the peak value of carbon dioxide in the low carbon scenario is up to 0.98 million tons 31294 in 2033, the peak value of carbon emissions in the high speed scenario is up to 0.37 million tons 30251 in 2036, and the peak value of carbon emissions in the baseline scenario is up to 0.61 million tons 26243 in 2038. Based on the peak time and prediction results of carbon emissions under the three scenarios, the main factors contributing to the reduction of carbon emissions in Guizhou Province are analyzed, and important data basis is provided for energy conservation and emission reduction in Guizhou Province.

## 1. Introduction

In recent years, the problem of global warming has attracted the attention of all countries in the world. A large number of greenhouse gas emissions are the main factors causing climate warming. How to reduce greenhouse gas emissions to alleviate global warming is a problem that people all over the world need to work together to solve. The United Nations Intergovernmental Panel on Climate Change (IPCC) Fifth Assessment of Climate Change Bulletin objectively stated that climate warming is mainly caused by the burning of large amounts of fossil fuels in the process of human activities, and alleviating global warming has become an unavoidable responsibility of all countries in the world. The global average temperature rise caused by excessive CO_2_ emissions seriously threatens the living space of human beings and the sustainable development of society.

Global climate change is closely related to the sustainable development of all countries in the world. In order to actively respond to global climate change, governments have taken relevant measures. China has put forward the ambitious goal of “achieving carbon peak by 2030 and achieving carbon neutrality by 2060”. In order to better implement this task, China has actively increased the research and development of new energy-related technologies to reduce the proportion of fossil energy use, thereby achieving a profound change in the energy consumption structure. In the aspect of protecting the ecological environment, we should make full use of the ecological environment resources such as water energy, wind energy and tidal energy, and carry out “pollution reduction and carbon reduction” in all aspects of ecological and environmental protection work. We should strengthen the guidance of typical demonstrations, fully mobilize the enthusiasm of local governments, departments, industries and enterprises, and jointly build a good working pattern.

As a major Karst Areas in China, the energy consumption of Guizhou Province is mainly coal, oil and other primary energy consumption. In recent years, the accelerated development of urbanization and rapid economic growth have made its dependence on traditional energy increasing, and the total energy consumption has been at a high level for a long time. Under the dual impetus of urbanization and economic development, the carbon emissions caused by the growth of energy consumption in Guizhou Province are increasing year by year. As a major energy province, the effect of energy saving and emission reduction is related to the overall situation of low-carbon economic development in China. In the process of realizing the peak of carbon emissions in China in 2030, it is particularly important to implement efficient emission reduction policies, so it is necessary to study the impact mechanism of carbon emissions in depth, monitor the level of carbon emissions with scientific and effective methods and predict them accurately. Based on different neural network models, the carbon peak value of Guizhou Province is predicted, which is of great significance for Guizhou Province to achieve the goal of carbon emission reduction and the grand goal of “3060” in China(Figure 1).

**Figure 1.**
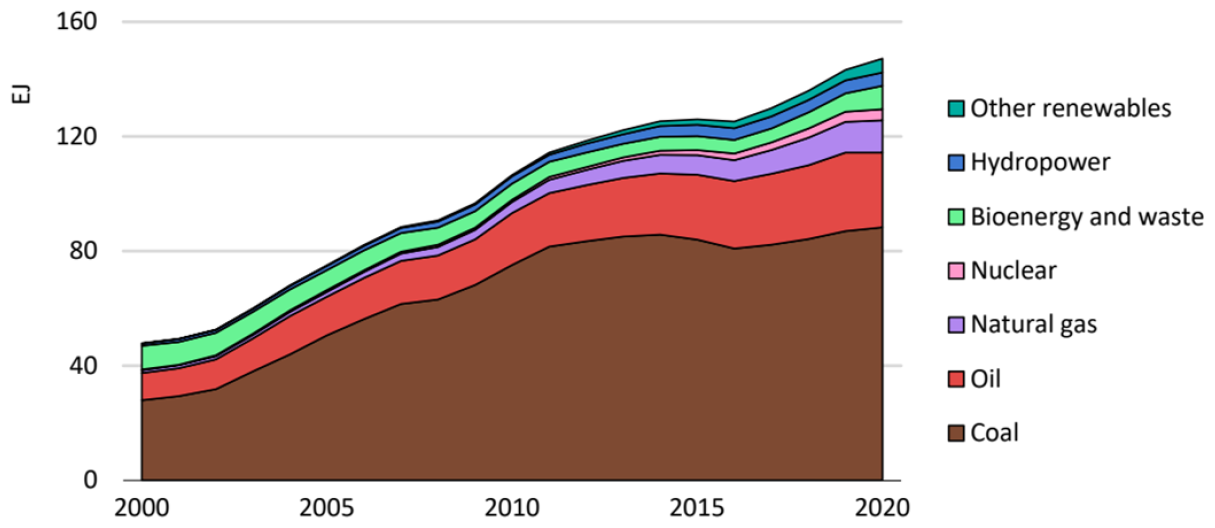
Total primary energy demand by fuel in China

The existing research on carbon emissions is mostly focused on the national or industry level, and there are few literatures on the regional level. In this paper, Guizhou Province, a key karst region in southwest China, is taken as the research object. Firstly, the total carbon emissions of Guizhou Province are calculated through the relevant data of energy consumption in Guizhou Province, and the current situation of carbon emissions in Guizhou Province is analyzed. Then, by reading relevant literature, the influencing factors of carbon emissions in Guizhou Province are determined, and by using machine learning technology, the influencing factors with high correlation are screened and the future trend of carbon emissions is predicted. Finally, the future carbon emissions in Guizhou Province are analyzed from the main factors affecting carbon emissions, the time of carbon peak and the peak value of carbon emissions. Therefore, this paper is of great significance to enrich the theoretical research on carbon emissions at the regional level.

## 2. Literature review

The national carbon emission reduction work needs to be jointly promoted by all regions of the country, and regions with different levels of industrial development should implement differentiated policies. As an important old industrial base in China, Guizhou’s traditional industries mainly use fossil energy as fuel, and its greenhouse gas emissions have been at a high level for a long time, which poses a more severe challenge for Guizhou to achieve carbon peak on schedule, so a more scientific and accurate calculation method is particularly important. Predicting the level of carbon emissions in Guizhou Province is of guiding significance for the implementation of energy conservation and emission reduction in Guizhou Province.Based on the optimized neural network model, this paper predicts and analyzes the peak carbon emissions in Guizhou Province in the next 20 years, which can clarify the existing problems and gaps, and help Guizhou Province to formulate the direction of energy development.

### 2.1. Research on influencing factors of carbon emission

The When studying the influencing factors of carbon emissions,scholars pay more attention to the contribution of different influencing factors to carbon emissions, and mainly select economic indicators such as population, economy and energy intensity to construct the index system of influencing factors of carbon emissions. Ang[1] analyzed the change of carbon dioxide produced per unit of electricity in the world, and took the import and export of each country, the fuel structure of power generation and emission factors as the main factors affecting carbon emissions. The study revealed that the improvement of power generation efficiency was the main reason for the reduction of CO_2_ emissions. Rustemoglu[2] studied the carbon dioxide levels of Brazil and Russia from 1992 to 2012, decomposed the factors affecting carbon emissions into employment, economy, carbon emission intensity, and found that Brazil’s carbon dioxide emissions were not decoupled from its economic development. Russia’s carbon dioxide emissions have been greatly reduced with the increase of energy intensity. Lin[3] used the input-output method to analyze the carbon emissions of the national food industry, and divided the industrial carbon emissions into four main factors: pollution factor, total output, energy intensity and energy structure. Roinioti[4] regards the development level of national economy, energy consumption intensity, fuel consumption capacity and carbon emission intensity as the main factors affecting carbon dioxide. Kim[5] decomposed the factors affecting carbon emissions into production scale and production intensity, and made corresponding research and analysis on the contribution of energy consumption growth in various sub-industries. Roman[6] took Colombia as the research object, and used the IDA-LMDI model to decompose the influencing factors of CO_2_ emissions into five aspects, including energy intensity, wealth value, fossil fuel substitution and renewable energy development, to explore the contribution level of CO_2_ increment.

Chinese scholars in the establishment of carbon emissions impact factors index system, mostly from the perspective of energy structure, population growth, economic and other factors. Liu Ying[7] and others take China’s steel industry as the main research object, and the results show that energy intensity has a greater impact on carbon emissions, while the consumption structure has not played the expected role. Dewey [8] deeply studied the influencing factors of carbon dioxide generated by indirect consumption in daily life of Chinese residents, and found that the socio-economic level is the main driving factor affecting the CO_2_ emissions of urban and rural residents, and the contribution of urban and rural structure and the proportion of urban and rural consumption are inversely related to CO_2_ emissions. Fu Jingyan[9] used regression analysis to study the carbon emissions of thermal power industry in Guangdong Province, China. Liu Ting[10] and others used the LMDI method to conclude that economic growth and carbon dioxide levels change in the same direction, that is, the faster the economic growth, the more obvious the effect of promoting carbon dioxide emissions, while the effective utilization and conversion of energy can reduce the level of carbon dioxide emissions. Ma Xiaoming[11] et al. Used the LMDI decomposition model to study the carbon emissions of 30 provinces and cities in China from 2004 to 2014, and explored the contribution of the factors affecting carbon emissions by dividing the time period with 2009 as the demarcation point. The results showed that the growth of gross national product had the greatest impact on the national carbon emissions, while the contribution of other factors was weak. Wang Ying[12] and others selected energy intensity, economic development and population size to study the influencing factors of carbon emissions. Bai Xiaoyong[13] used high-resolution spatial data to put carbonate chemical weathering carbon sink, silicate chemical weathering carbon sink, vegetation-soil ecosystem carbon sink and energy carbon emissions on the spatial grid. Then a carbon neutral index model was established to reveal the contribution of terrestrial ecosystem carbon sink to carbon neutrality. Finally, the results of the study are compared with those of other countries from the horizontal and vertical perspectives. This study provides a new research idea for the measurement of carbon neutralization capacity, and provides an important reference value and data basis for the systematic determination of global carbon neutralization capacity.the model made by Bai Xiaoyong’s team is highly recognized in the academic circles.(Figure 2).

**Figure 2.**
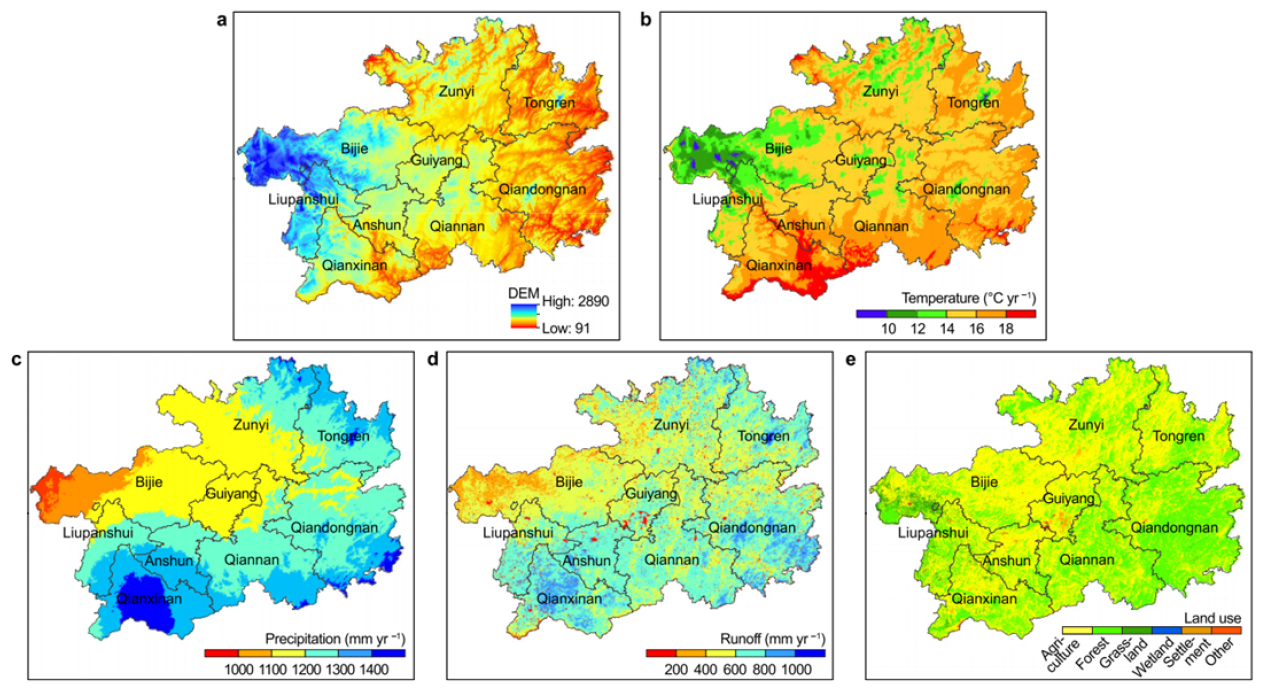
The spatial pattern of digital elevation model

### 2.2. Domestic and foreign carbon emission prediction research

Existing studies mainly predict the carbon emissions of different countries or a certain industry through logistic regression model, STIRPAT model, scenario analysis and other methods. Ouedraogo [14] uses the LEAP framework to model the analysis and projections of energy demand and associated emissions under alternative strategies in Africa from 2010 to 2040. Lin [15] et al. Conducted a survey on China’s manufacturing industry based on the STIRPAT model, and found that macroeconomic growth factors determine the carbon dioxide emissions of the industry, while the effects of fuel utilization rate and urbanization rate have significant regional heterogeneity. Wang [16] et al. Constructed STIRPAT model to study the influencing factors of carbon emissions in Xinjiang from 1952 to 2012, and found that there were differences in the impact of various factors on carbon emissions in different historical periods. Before 1978, the expansion of population size was the main factor causing the increase of carbon emissions. From 1978 to 2000, economic growth and population size are the main driving factors for the rise of carbon emissions, and after 2000, the main factors for the increase of carbon emissions are economic development and fixed asset investment. Kachoee [17] and others used the LEAP model to predict the carbon dioxide emissions of Iran’s power sector in the next 30 years, and concluded that the level of economic growth is the main factor affecting carbon dioxide emissions(Figure 3). Emodi[18] and others used the LEAP model to study climate change in Australia’s power sector, and found that reducing expenditure on environmental protection and resource conservation would produce economic benefits.

**Figure 3.**
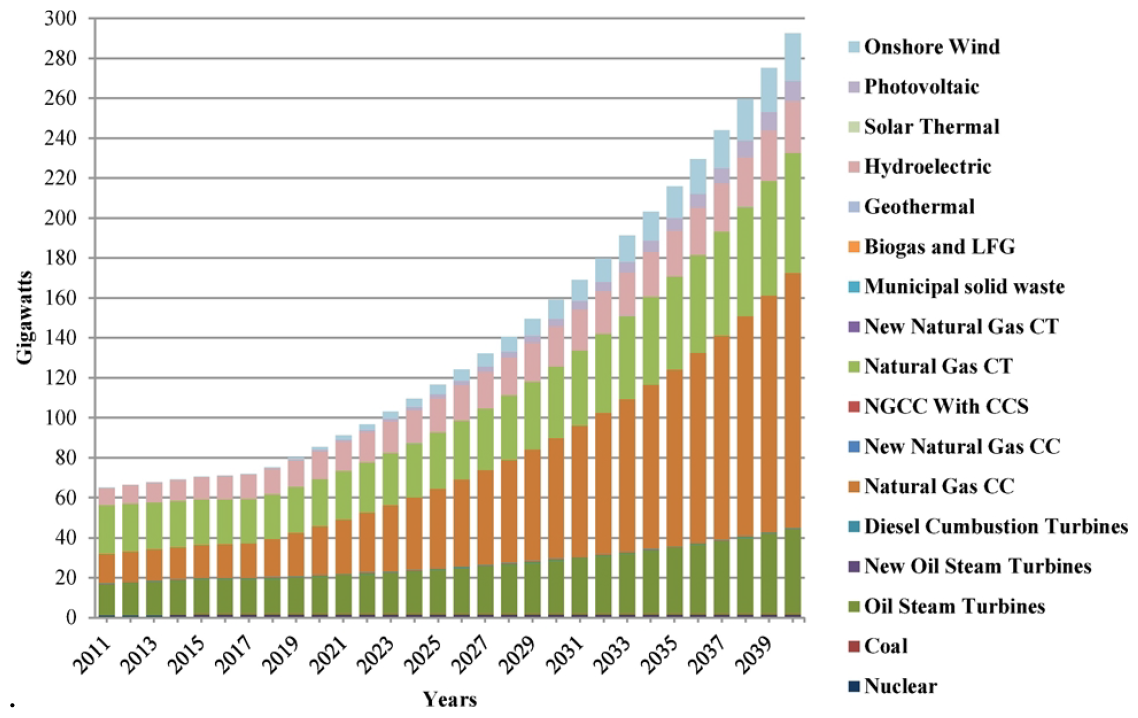
Generation capacity in the BAU scenario until 2040

### 2.3. Research on neural network model

At present, neural network model has been widely used in different fields. The representative neural network models are BP neural network, radial basis function network, Hopfield model, GMDH network, adaptive resonance theory, Boltzmann machine, CPN model and so on. Lapedes et al. First used neural networks for economic forecasting in 1987. Wang Chunjuan[27] mainly applied neural network to the prediction of typhoon, debris flow and geological subsidence. Fan Decheng[28] and others established POS-BP neural network model to predict the total carbon emissions and intensity of 30 provinces, municipalities and autonomous regions in China. Liu Ying [29]compared the advantages of neural network and other traditional prediction methods, used BP neural network model combined with terminal information collection system and Web Service technology to design the intelligent system of urban road-occupying parking, and proved the feasibility of the management system through actual data. Liu Xiaowei [30] predicted the price of stock investment by combining neural network model with principal component analysis and multiple linear regression. Huo Xiaolong [31] and others studied the problem of gas outburst in tunnels, and used BP neural network to predict it, and achieved good results, verifying that the predicted value and the real value have good consistency. Tang Xiaocheng [32] used BP neural network to study the prediction of air pollutant concentration. The original BP neural network was used to calculate the system error of all samples through successive iterations and batch processing, which improved the execution efficiency.

To sum up, in the research on the influencing factors of carbon emissions, scholars mainly take population, economy, energy structure and other factors as the influencing factors of carbon emissions to establish the carbon emission index system, and apply it to the follow-up prediction research. In the study of carbon emissions forecasting, scholars mainly use traditional econometric methods such as logistic regression model, STIRPAT model, scenario analysis and so on. The neural network model has achieved good results in economic forecasting. By reviewing the carbon emissions related literature at home and abroad, it is found that most of the current research focuses on the national level or industry level, and there are few studies on the use of machine learning algorithms to predict the peak carbon emissions, and the neural network model used is relatively single. The training of single neural network prediction model takes a long time and is easy to fall into local optimum. Therefore, this paper combines neural network model with carbon emissions research closely, and optimizes different algorithms to carry out the prediction of carbon emissions in Guizhou Province.

## 3. Methods

In this paper, the carbon emissions of Guizhou Province are calculated by the inventory method based on the energy consumption data of Guizhou Province. Referring to the relevant literature and combining with the actual development of Guizhou Province, 12 factors affecting the carbon emissions of Guizhou Province are selected to establish the characteristic subset. By introducing the grey correlation analysis method, the indicators with higher influence degree are selected and applied to the follow-up prediction research. Finally, the carbon emissions from 2020 to 2040 in Guizhou Province are predicted under three different development scenarios by establishing a prediction model based on WOA-ELM.

### 3.1. Calculation of Carbon Emissions in Guizhou Province

Because there is no authoritative agency in China to directly provide the test data of carbon dioxide emissions, this paper uses a compromise method to discount the emission data of each year in different collected data and take the average value. The calculation of carbon emissions and the collection of relevant data will vary depending on the subject of the study. In view of the characteristics of carbon emissions in Guizhou Province and the difficulty of data acquisition, this paper uses the method of estimating carbon emissions proposed by the United Nations Intergovernmental Panel on Climate Change (IPCC) in 2006. The method is based on the gas emission inventory, and calculates the carbon emission generated by energy consumption by multiplying the energy consumption data by the energy carbon emission factor. The specific formula is as follows:.

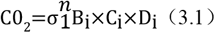

Where B_i_ represents the consumption of the i-th energy source, and n represents the type of energy source. In this paper, n=9 represents coal, coke, crude oil, gasoline, kerosene, diesel oil, natural gas and electricity in energy consumption. The conversion coefficient and carbon emission coefficient of each energy standard coal in Guizhou Province are shown in Table 1:

**Table 1.**
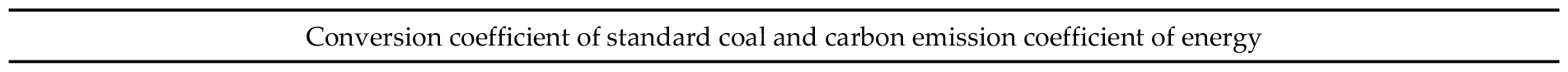
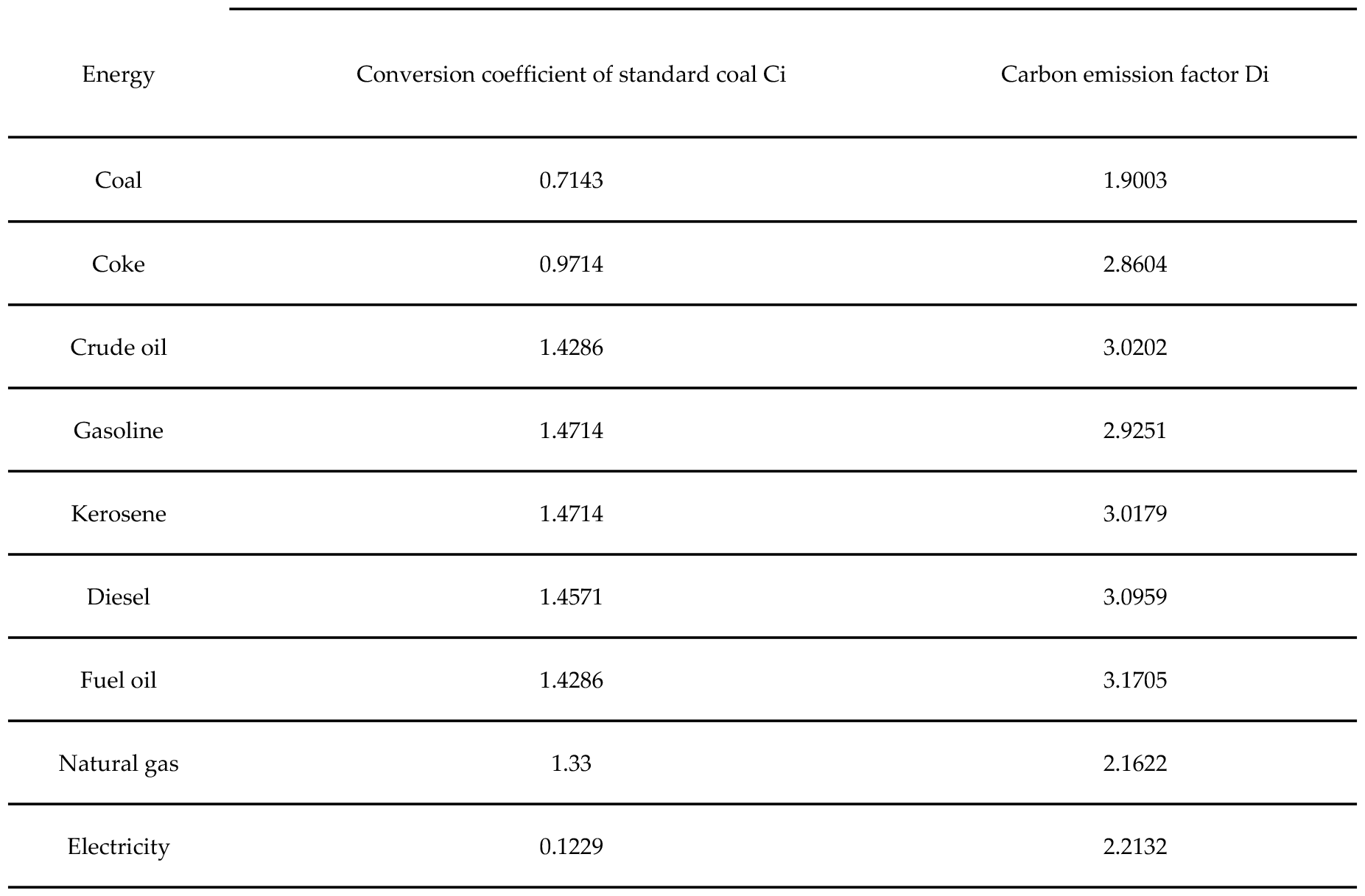
Standard Coal Conversion Coefficient and Energy Carbon Emission Coefficient.

| Energy | Conversion coefficient of standard coal Ci | Carbon emission factor Di |
| --- | --- | --- |
| Coal | 0.7143 | 1.9003 |
| Coke | 0.9714 | 2.8604 |
| Crude oil | 1.4286 | 3.0202 |
| Gasoline | 1.4714 | 2.9251 |
| Kerosene | 1.4714 | 3.0179 |
| Diesel | 1.4571 | 3.0959 |
| Fuel oil | 1.4286 | 3.1705 |
| Natural gas | 1.33 | 2.1622 |
| Electricity | 0.1229 | 2.2132 |

This section may be divided by subheadings. It should provide a concise and precise description of the experimental results, their interpretation, as well as the experimental conclusions that can be drawn.

It can be seen from Table 2 that from 2000 to 2012, the total carbon emissions of Guizhou Province were on the rise. From 2012 to 2016, the total carbon emissions in Guizhou Province showed a downward trend, which is related to the construction of ecological civilization in Guizhou Province during this period, and the practice of green mountains is Jinshan and Yinshan. In 2020, the total amount of carbon emissions in Guizhou Province was 1.1 million tons 22237, about twice as much as in 2002. With the development of social economy, the carbon emissions in Guizhou Province will show a steady upward trend in the future.However, due to the inhibition of carbon emissions by the implementation of policies such as the introduction of carbon emission reduction policies, the establishment of carbon trading market and the increase of the proportion of new energy applications in Guizhou Province, the growth rate of carbon emissions will gradually decrease, and the development trend will slow down year by year.

**Table 2.**
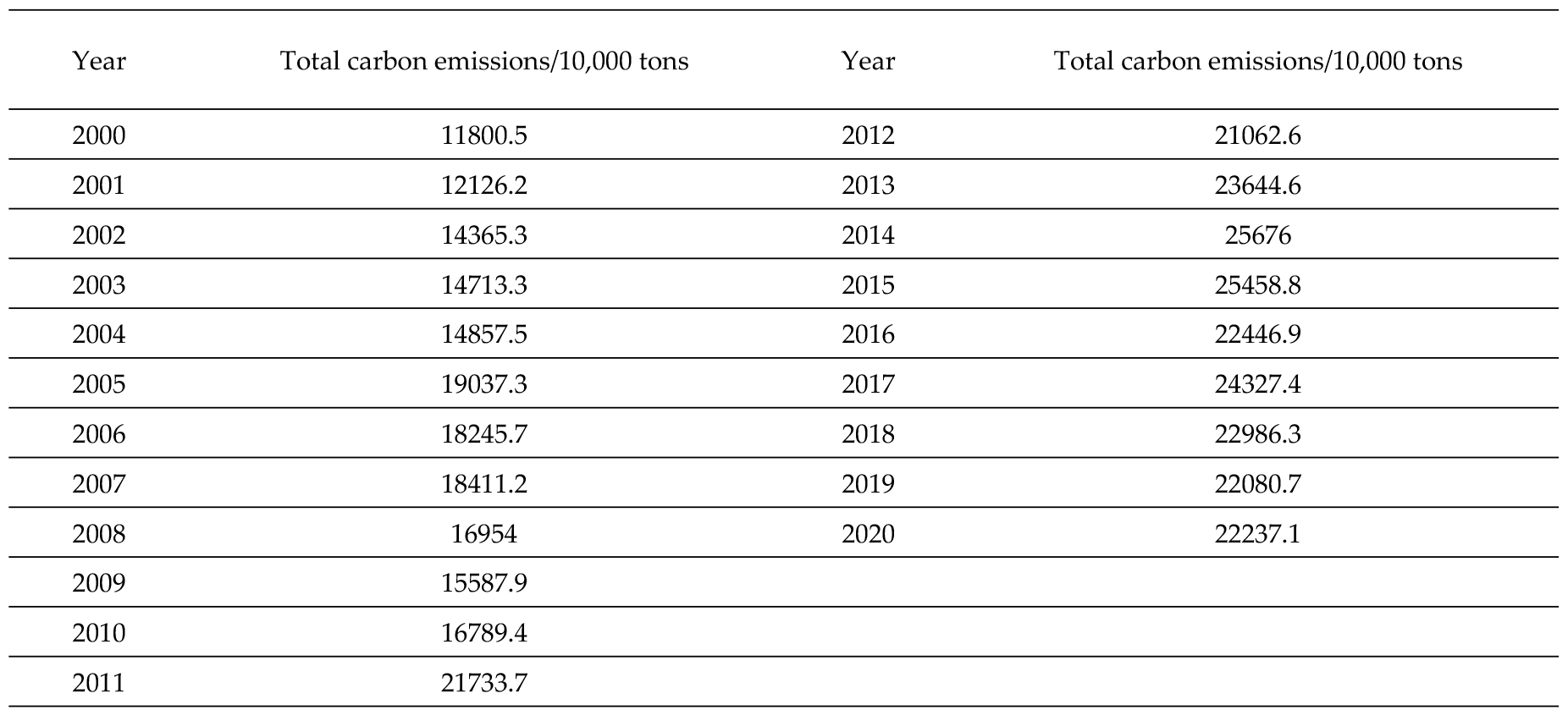
Carbon emission of Guizhou Province from 2000 to 2020 (converted value)

| Year | Total carbon emissions/10,000 tons | Year | Total carbon emissions/10,000 tons |
| --- | --- | --- | --- |
| 2000 | 11800.5 | 2012 | 21062.6 |
| 2001 | 12126.2 | 2013 | 23644.6 |
| 2002 | 14365.3 | 2014 | 25676 |
| 2003 | 14713.3 | 2015 | 25458.8 |
| 2004 | 14857.5 | 2016 | 22446.9 |
| 2005 | 19037.3 | 2017 | 24327.4 |
| 2006 | 18245.7 | 2018 | 22986.3 |
| 2007 | 18411.2 | 2019 | 22080.7 |
| 2008 | 16954 | 2020 | 22237.1 |
| 2009 | 15587.9 |  |  |
| 2010 | 16789.4 |  |  |
| 2011 | 21733.7 |  |  |

### 3.2. Selection of Influencing Factors of Carbon Emission in Guizhou Province

By summarizing and synthesizing the relevant literature on the influencing factors of carbon emissions, it is found that most scholars usually choose the influencing factors of carbon emissions from the diversified perspectives of macro-economy, industrial structure, energy consumption and scientific and technological development. According to the actual social and economic development of Guizhou Province, this paper comprehensively considers the scientific principle, systematic principle and authenticity principle of index selection. Total population, urbanization rate, household consumption level, per capita GDP, energy intensity, carbon emission intensity, foreign direct investment, energy structure, the proportion of primary industry, the proportion of secondary industry, the proportion of tertiary industry, total energy consumption, a total of 12 factors are selected and qualitatively analyzed.

1. Total population. With the expansion of population size and the continuous improvement of living standards, people’s demand for food, clothing, shelter and living environment in daily life will increase, which will drive the development of related industries, and the energy demand will increase, which will lead to the increase of CO_2_ emissions.
2. Urbanization rate. Urbanization rate is used to reflect the progress of urbanization in a country or region. The improvement of urbanization rate can drive the rise of economic development level, thus promoting the process of local industrialization. The further promotion of industrialization in Guizhou Province will inevitably generate a large amount of energy demand. If we only pay attention to the development of urbanization and ignore the resources and environment, it will inevitably lead to a large amount of energy consumption and the vicious growth of carbon dioxide.
3. Residents’ consumption level. Due to the rising consumption level of residents, the pace of urban industrialization and rapid economic growth have been accelerated, but the total carbon emissions have increased significantly due to the energy consumption.
4. GDP per capita. Generally speaking, per capita GDP reflects a country’s economic development and people’s living standards, but at the same time, it can also express the growth level of carbon emissions caused by the continuous expansion of economic development and scale to a certain extent. Economic and social development can not be separated from industrial development, and industrial development can not be separated from energy power. The rapid development of global economy will inevitably lead to a large amount of energy consumption, resulting in a large amount of carbon dioxide emissions.
5. Energy intensity. The amount of energy consumed per unit of output reflects the dependence of social and economic development on energy. Lower energy intensity means lower energy use, higher energy efficiency, and increased output [46].
6. Carbon emission intensity. Carbon emission intensity refers to the proportional relationship between total carbon emissions and GDP, which shows the relationship between social and economic development of a country or region and carbon emissions. Generally speaking, the better the economic and social development, the higher the level of industrial development, and the lower the carbon emission intensity [47]. With the development of economy and the progress of science and technology, the intensity of carbon emissions will gradually decrease.
7. Foreign direct investment. Since the reform and opening up, the speed of introducing and utilizing foreign direct investment in Guizhou Province has been increasing. However, with the economic development, environmental problems have become more severe. Considering the negative impact of foreign direct investment on carbon emissions and other environmental problems in the region, it is very necessary to study the impact of foreign direct investment on carbon emissions in Guizhou Province.
8. Energy structure. The proportion of coal consumption has an important impact on total carbon emissions [49]. Set the proportion of coal energy consumption and fossil energy consumption to represent the energy structure.
9. Proportion of primary industry. The primary industry is a production sector with agriculture as its main object and nature as its direct object. The higher the proportion of the primary industry, the lower the degree of industrial modernization in the region. In the process of continuous economic development, the proportion of the primary industry will gradually decrease in the proportion of the industrial structure.
10. Proportion of secondary industry. The secondary industry is represented by industrial enterprises. There are many sub-industries with high energy consumption and high pollution. Many links promote the increase of energy consumption, resulting in a large amount of carbon dioxide emissions [50]. Selecting the proportion of secondary industry can better show the level of industrial structure in Guizhou Province.
11. Proportion of tertiary industry. As an industrialized city, the proportion of tertiary industry in Guizhou Province has gradually increased in recent years, and the rational transformation of industrial structure will promote low-carbon technological innovation and have a significant positive impact on reducing carbon emissions.
12. Total energy consumption. Energy is an important material basis for economic development, and energy consumption plays a decisive role in economic growth. Different energy structures have different degrees of environmental pollution [51].

The selected indicators are shown in Table 3.

**Table 3.** Influencing Factors and Indicators of Carbon Emission in Guizhou Province.

| Indicator layer | Unit |
| --- | --- |
| Total population | Ten thousand people |
| Urbanization rate | % |
| Residents' consumption level | yuan/person |
| GDP per capita | yuan/person |
| Energy intensity | Ton of standard coal/10,000 yuan |
| Carbon emission intensity | Ton/10,000 yuan |
| Foreign direct investment | Ten thousand dollars |
| Energy structure | % |
| Proportion of primary industry | % |
| The proportion of the secondary industry | % |
| Proportion of tertiary industry | % |
| Total energy consumption | Ten thousand tons of standard coal |

## 4. Results

The total carbon emissions of Guizhou Province from 2000 to 2020 are calculated as the basic data, and the population, economy and energy data of Guizhou Province from 2000 to 2020 are selected to calculate the correlation between the total carbon emissions of Guizhou Province and its influencing factors according to the above method. The data used are from the Statistical Yearbook of Guizhou Province 2000-2020.

In order to facilitate the subsequent modeling and data representation, the variable names are redefined. X1, X2, X3, X4, X5, X6, X7, X8, X9, X10, X11 and X12 are used as independent variables to represent 12 factors affecting carbon emissions in Guizhou Province.The total carbon emission Y of Guizhou Province is set as the reference sequence, and the 12 influencing factors are set as the comparison sequence. The original data is normalized to eliminate the dimensional influence. Table 4 shows the results of data normalization.

**Table 4.** Normalization results of indicator data.

| Year | Y | X1 | X2 | X3 | X4 | X5 | X6 | X7 | X8 | X9 | X10 | X11 | X12 |
| --- | --- | --- | --- | --- | --- | --- | --- | --- | --- | --- | --- | --- | --- |
| 2000 | 0 | 0 | 0 | 0 | 0 | 1 | 0 | 1 | 0.65 | 0.04 | 0.65 | 0.09 | 0.04 |
| 2001 | 0.04 | 0.08 | 0.01 | 0.02 | 0.01 | 0.86 | 0.01 | 0.94 | 0.59 | 0.04 | 0.59 | 0.1 | 0.04 |
| 2002 | 0.08 | 0.16 | 0.03 | 0.03 | 0.03 | 0.7 | 0.03 | 0.93 | 0.59 | 0.03 | 0.59 | 0.2 | 0.03 |
| 2003 | 0.06 | 0.23 | 0.04 | 0.04 | 0.04 | 0.59 | 0.03 | 0.92 | 0.54 | 0 | 0.54 | 0.22 | 0 |
| 2004 | 0.09 | 0.28 | 0.05 | 0.06 | 0.05 | 0.55 | 0.02 | 0.66 | 0.55 | 0.02 | 0.55 | 0.27 | 0.02 |
| 2005 | 0.22 | 0.32 | 0.06 | 0.07 | 0.07 | 0.56 | 0.04 | 0.64 | 0.68 | 0.11 | 0.68 | 0.24 | 0.11 |
| 2006 | 0.17 | 0.36 | 0.07 | 0.09 | 0.09 | 0.5 | 0.06 | 0.4 | 0.58 | 0.11 | 0.58 | 0.34 | 0.11 |
| 2007 | 0.23 | 0.39 | 0.09 | 0.1 | 0.1 | 0.44 | 0.09 | 0.56 | 0.54 | 0.1 | 0.54 | 0.37 | 0.1 |
| 2008 | 0.31 | 0.41 | 0.1 | 0.1 | 0.13 | 0.41 | 0.15 | 0.8 | 0.57 | 0.15 | 0.57 | 0.37 | 0.15 |
| 2009 | 0.42 | 0.44 | 0.13 | 0.12 | 0.15 | 0.44 | 0.14 | 0.72 | 0.43 | 0.28 | 0.43 | 0.41 | 0.28 |
| 2010 | 0.55 | 0.45 | 0.55 | 0.16 | 0.21 | 0.34 | 0.08 | 0.82 | 0.56 | 0.31 | 0.56 | 0.35 | 0.31 |
| 2011 | 0.61 | 0.62 | 0.56 | 0.18 | 0.26 | 0.31 | 0.17 | 0.77 | 0.61 | 0.41 | 0.61 | 0.34 | 0.41 |
| 2012 | 0.59 | 0.72 | 0.57 | 0.23 | 0.33 | 0.27 | 0.28 | 0.59 | 0.65 | 0.52 | 0.65 | 0.31 | 0.52 |
| 2013 | 0.67 | 0.78 | 0.6 | 0.31 | 0.43 | 0.2 | 0.38 | 0.45 | 1 | 0.6 | 1 | 0 | 0.6 |
| 2014 | 0.73 | 0.87 | 0.61 | 0.36 | 0.48 | 0.19 | 0.51 | 0.33 | 0.78 | 0.69 | 0.78 | 0.23 | 0.69 |
| 2015 | 0.85 | 0.99 | 0.68 | 0.46 | 0.61 | 0.14 | 0.7 | 0.17 | 0.9 | 0.82 | 0.9 | 0.14 | 0.82 |
| 2016 | 0.94 | 1 | 0.76 | 0.58 | 0.75 | 0.1 | 0.83 | 0.11 | 0.94 | 0.94 | 0.94 | 0.12 | 0.94 |
| 2017 | 1 | 0.99 | 0.82 | 0.69 | 0.85 | 0.07 | 0.92 | 0 | 0.85 | 1 | 0.85 | 0.2 | 1 |
| 2018 | 0.96 | 0.95 | 0.85 | 0.78 | 0.94 | 0.01 | 1 | 0.31 | 0.82 | 0.87 | 0.82 | 0.23 | 0.87 |
| 2019 | 0.93 | 0.93 | 0.88 | 0.88 | 1 | 0 | 0.94 | 0.26 | 0.68 | 0.88 | 0.68 | 0.39 | 0.88 |
| 2020 | 0.87 | 0.86 | 0.91 | 0.95 | 1 | 0 | 0.14 | 0.13 | 0.4 | 0.87 | 0.4 | 0.63 | 0.87 |

Calculate the difference between the maximum and minimum absolute values in the matrix, set the resolution coefficient as 0.5 to obtain the correlation coefficient table, and take the average value of the correlation coefficients of different sequences at each time to obtain the correlation degree and sort them. The results are shown in Table 5.

**Table 5.** Correlation Degree Ranking of Influencing Factors.

| Influencing factors | Correlation degree X | Incidence order |
| --- | --- | --- |
| X1 | 0.8573 | 2 |
| X2 | 0.8534 | 3 |
| X3 | 0.7589 | 5 |
| X4 | 0.7953 | 4 |
| X5 | 0.5046 | 12 |
| X6 | 0.4870 | 11 |
| X7 | 0.6907 | 6 |
| X8 | 0.5060 | 10 |
| X9 | 0.5166 | 9 |
| X10 | 0.6632 | 8 |
| X11 | 0.6968 | 7 |
| X12 | 0.8801 | 1 |

The closer the correlation degree is to 1, the stronger the correlation degree between the influencing factors and carbon emissions in Guizhou Province is. It can be seen from the above table that the top five correlation degrees are X12, X1, X2, X4 and X3. The influencing factors they represent are total energy consumption, total population, urbanization rate, per capita GDP and residents’ consumption level. The correlation degrees are all greater than 0.75, which are the main correlation factors. Secondly, the correlation degree of X7, X11, X10, X9, X8 and X5 is between 0.5 and 0.7, which belongs to the medium correlation degree. The correlation between energy intensity and carbon emissions in Guizhou Province is low, and the correlation degree is 0. 487. collects the relevant data of energy consumption in Guizhou Province, calculates and analyzes the carbon emissions in Guizhou Province from 2000 to 2020. The results show that the total carbon emissions in Guizhou Province are closely related to the economic development situation and relevant policies. Referring to the existing literature and combining with the actual situation of Guizhou Province, this paper initially sets twelve indicators, such as the total population, urbanization rate, consumption level of residents, per capita GDP, as the influencing factors of carbon emissions in Guizhou Province, and expounds the reasons for the selection in detail. It can be known that the total energy consumption, the total population, the urbanization rate, the per capita GDP and the residents’ consumption level have a high correlation with the carbon emissions in Guizhou Province, and the five factors with strong correlation can be used as the input variables of the prediction model to improve the accuracy of predicting the carbon emissions in Guizhou Province.

## 5. the design of carbon emission prediction model and carbon peak prediction

### 5.1. Design of prediction model

#### 5.1.1 Establishment of BP neural network model

BP neural network is composed of input layer, hidden layer and output layer. In the process of establishing BP neural network, the number of hidden layer nodes has a very important impact on the fitting effect of the model. The current research does not define how to set more accurate hidden layer nodes. Setting too many or too few hidden layer nodes will affect the results of the data: too many hidden layers are prone to over-fitting, resulting in increased training time, and too few hidden layer nodes will affect the accuracy of the data fitting. The number of hidden layers needs to be determined according to the characteristics of the data. In this paper, the number of hidden layer nodes [53] will be determined by trial and error. The specific settings of each level and node are as follows:

##### (1) Setting of model input layer and output layer

The five factors selected in the third chapter are the main factors affecting the carbon emissions of Guizhou Province, so these data are used as the input variables of the model, that is, the nodes of the input layer, including the total population, urbanization rate, household consumption level, per capita GDP, and total energy consumption. The number of neurons in the output layer of the model is 1, that is, the carbon emission of Guizhou Province.

##### (2) Analysis of network structure parameters

###### 1) Selection of network layer number

BP neural network is a multilayer neural network, and the selection of the number of layers is very important for establishing a reasonable network model. Different number of hidden layers will affect the final effect of model fitting. When the number of hidden layers is increased, the network structure becomes complex, the prediction ability of the model is improved, and the error is reduced, but the overfitting phenomenon is prone to occur, and the training time is not ideal. Relatively, the number of hidden layers is reduced, the network structure is simple, the prediction ability of the model will be reduced, the accuracy will be reduced, the error will be increased, but the training speed is fast. Hornik proved that the three-layer BP network structure can meet the fitting requirements of most nonlinear systems, as long as the number of hidden layer nodes is set appropriately, it can also help the network to fit most functions with high accuracy. Therefore, on the basis of meeting the needs of training accuracy, this paper chooses a three-layer BP neural network with only one hidden layer to predict the carbon emission data.

###### 2) Determination of activation function and other parameters

The function of activation function is to introduce the nonlinear relationship into neurons through mapping. In order to better represent the nonlinear relationship of the function, it is necessary to select the appropriate type of activation function. The hyperbolic tangent function is a common activation function, which maps the number taking the value of (-∞, + ∞) into (−1,1), so that the variable is in the largest possible threshold range, which can better preserve the nonlinear variation level of the function. Therefore, the transfer function of the hidden layer node is chosen as the tangent S-type transfer function tansig in this paper. The input and output values of the linear transfer function purelin can take any value. In order to facilitate the comparison with the sample value, purelin is selected as the output value returned by the output layer.Learning rate refers to the change of information accumulation speed of BP neural network with time. Different learning rate settings will affect the training time and training effect of the model. The training speed of the model with a larger learning rate value will be relatively fast, but there will be large fluctuations in the later period, resulting in the model can not converge. When its value is small, although it can make the simulation results of the model more accurate, it will greatly increase the training time. In general studies, the learning rate γ is usually set between (0,1). In this paper, through continuous debugging and comparison of the training effect in the training process, the learning rate γ = 0.1 is finally selected. The accuracy of network training is required to be 0.001, and the maximum number of training times is 500.

###### 3) Selection of the number of neurons in the hidden layer

The number of neurons in the hidden layer affects the performance of the network. The more the hidden layer nodes are, the more complex the model training is, and the more time-consuming it is, and it may not be able to achieve function approximation smoothly, resulting in a shock effect. If the number of neurons in the hidden layer is too small, the training accuracy of the network may not meet the ideal requirements, resulting in the failure of the experimental results. After defining the relevant network parameters and activation functions, the approximate range is calculated by empirical formula, and then the number of hidden layer neurons is determined by debugging single variable trial and error. According to the existing empirical formula, there are the following three quantitative relationships:

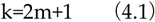

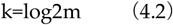

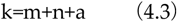

Where K is the number of nodes in the hidden layer, m is the number of nodes in the input layer, n is the number of nodes in the output layer, and a is an arbitrary constant between 1 and 10. This paper uses the empirical formula to calculate that the number of nodes in the hidden layer can be selected between 4 and 13. After adjusting the parameter settings for many times, combined with the overall level of the training results, as shown in Table 6, the number of nodes in the hidden layer is set to 10. Finally, it is determined that the number of neurons in the input layer of the BP neural network used in this paper is 5, the number of nodes in the hidden layer is 10, the number of nodes in the input layer is 1, the learning rate is 0.1, and the setting accuracy is 0.001.

**Table 6.** Model effect of different hidden layer node number.

| Implies the number of layer nodes | Number of training sessions | Mean absolute percent error |
| --- | --- | --- |
| 4 | 200 | 7.25 |
| 5 | 102 | 8.04 |
| 6 | 133 | 6.36 |
| 7 | 56 | 10.04 |
| 8 | 74 | 9.28 |
| 9 | 120 | 6.03 |
| 10 | 42 | 4.37 |
| 11 | 86 | 5.41 |
| 12 | 67 | 6.79 |
| 13 | 50 | 11.26 |

#### 5.1.2. Establishing Extreme Learning Machine Model

The steps of the algorithm can be divided into:

Step 1, firstly determining the number of neurons of a hidden layer, and randomly generating a connection value between an input layer and the hidden layer and a neuron bias bi of the hidden layer in a network model;

Step 2, determining an activation function of the neurons of the hidden layer, and calculating an output matrix H of the hidden layer by selecting an infinitely differentiable function;

Step 3: Finally calculate the output layer weight β ∗= H + T.

#### 5.1.3. Establishment of WOA-BP model

BP algorithm is a common learning algorithm in various fields, but the existing problems restrict its development. In the training process, the initial weights and thresholds are randomly generated, so that the generalization ability can not be well guaranteed, so WOA is used to optimize the initial parameters of BP neural network, so as to obtain a more stable WOA-BP neural network.

Steps of WOA optimizing BP neural network:

1. determine a BP neural network structure and initializing a weight value and a threshold value;
2. calculate that individual fitness of the whale, and taking a fitness function as an optimized target function;
3. set an algorithm stopping criterion, selecte different mechanisms, updating that individual positions of the whale, and optimizing parameters; 4, fin out that optimal whale position, and assigning the optimal weight value and the optimal threshold value to the BP neural network;

And step 5, training the optimized BP neural network and carrying out simulation test, and comparing the prediction performance of the BP neural network before optimization.

#### 5.1.4. Establishment of WOA-ELM model

Based on the whale optimization algorithm and the structure of extreme learning machine mentioned above, the WOA-ELM combination forecasting model is established. In the WOA algorithm, the optimal position of the humpback whale is the optimized ELM parameter value, and the WOA iteration is used to find the optimal wi and bi of the ELM, which can improve the prediction accuracy of the model.

(1) Initialize the ELM. The number of input neurons (set as 5), the number of hidden layer neurons (set as 10), the activation function type G (set as Sigmoid function), the input weight wi, and the hidden layer threshold bi.
(2) Initialize the parameters of WOA. Set the population size N (set to 50) and the maximum generation number Tmax (set to 500).
(3) initializing the position vector of the individual whale, connecting the randomly generated connection weight wi and nerve during the ELM training, The meta-bias bi is taken as the initial position vector of the individual whale. And (4) setting a fitness function (error rate in the ELM test), and calculating the current individual fitness in the initialized population to obtain the optimal (best fitness) whale individual.
(5) After the p value is randomly generated (0,1), the updated formula (Graph) (Graph) at different positions is determined by the values of A and p. When A (Graph) (Graph) values are different, there are three corresponding update position formula selections as follows: when A (Graph) (Graph)> 1, select to perform global random search; When A (Graph) (Graph)< 1, the random variable p-value is combined to choose between the shrinking enclosure and the spiraling strategy. And (6) judging whether the algorithm can reach a preset termination condition. When the end condition of the algorithm is reached, the algorithm is terminated, and the whale individual position vector with the best fitness value is output, that is, the optimal weight and threshold of the ELM network are obtained. Otherwise, the number of iterations is increased by one, and the step (5) is returned. And (7) inputting the finally obtained optimal parameters of the ELM network into the WOA-ELM model to predict the carbon emission of Guizhou Province.

### 5.2. Analysis and comparison of experimental results of prediction model

Because there is no authoritative agency in China to directly provide the test data of carbon dioxide emissions, this paper uses a compromise method to discount the emission data of each year in different collected data and take the average value. The calculation of carbon emissions and the collection of relevant data will vary depending on the subject of the study. In view of the characteristics of carbon emissions in Guizhou Province and the difficulty of data acquisition, this paper uses the method of estimating

#### 5.2.1. Prediction based on BP neural network model

##### (1) Simulation settings

The research data of this paper comes from the Statistical Yearbook of Guizhou Province from 2000 to 2020, China Energy Statistics Yearbook and the website of the National Bureau of Statistics. Through the third chapter of the calculation of carbon emissions data in Guizhou Province, for example, to verify the BP neural network prediction model of carbon emissions in Guizhou Province, the fitting degree of the evaluation model advantages and disadvantages.

##### (2) Prediction results and analysis

When dividing the training set and the test set, the relevant data from 2015 to 2020 are used as the training set data, and the remaining 6 groups are used as the test set data. Because the carbon emissions are changing year by year, in order to improve the prediction performance of the model, this paper predicts the carbon emissions year by year, and adds the influencing factors and carbon emissions data of the new year prediction to the training sample. Relative error and absolute error are used to analyze the prediction effect of the prediction model. Table 4.2 shows the prediction error of the test sample.

It can be seen from the carbon emission prediction curve in Table 7 that the change rule of the predicted value of carbon emission in Guizhou Province is generally consistent with the change rule of the real value, but the difference between the values is large, and the prediction effect is not stable enough. Among them, the difference between the predicted value and the real value in 2018, 2019 and 2020 is small, and the relative error is below 2.5%, which can meet our expectations for the prediction accuracy. However, there is a big difference between the predicted value and the real value in 2015, 2016 and 2017, which does not achieve the expected prediction effect. The main reason is that the initial weights and thresholds in BP neural network are determined randomly, and the parameters are difficult to achieve a good fitting in the model training, resulting in large fluctuations in the prediction results and unable to achieve a good prediction effect.

**Table 7.** Prediction error of BP neural network prediction model

| Year | Actual value/10,000t | Predicted value/10,000 tons | Relative error (%) | Absolute error/10,000 ton |
| --- | --- | --- | --- | --- |
| 2015 | 25458.8 | 22667.6 | 10.96 | 2791.2 |
| 2016 | 22446.9 | 22913.9 | -2.08 | -467 |
| 2017 | 24327.4 | 24815.2 | -2.01 | -487.8 |
| 2018 | 22986.3 | 22932.2 | 0.24 | 54.1 |
| 2019 | 22080.7 | 22457.6 | -1.71 | -376.9 |
| 2020 | 22237.1 | 21234.3 | 4.51 | 1002.8 |

#### 5.2.2. Prediction based on extreme learning machine model

The extreme learning machine model is used to predict the carbon emissions of Guizhou Province, the data from 2000 to 2014 is used as the training set, the data from 2015 to 2020 is divided into the test set, the training sample method is the same as the BP neural network structure, the relative error is reserved for two digits, the absolute error is reserved for one digit, and the error comparison between the actual value and the predicted value is obtained.

It can be seen from the prediction results in Table 8 that the model fits the carbon emission of Guizhou Province well and can approximate the relationship between the influencing factors and the carbon emission of Guizhou Province. However, the prediction results are unstable. Although the predicted values of most years are close to the actual values, the carbon emission obtained in 2017 is quite different from the actual values. In the forecast results, the absolute error between the forecast value and the real value in 2019 is 8.8, and the forecast value in this year is the closest to the actual carbon emissions of Guizhou Province in 2019. The average relative error of the test set is 0.43%, less than 5%. Compared with BP neural network, the prediction model of carbon emissions in Guizhou Province based on extreme learning machine has higher accuracy and stronger ability to approximate the nonlinear relationship of samples, but the setting of random initialization value and β also affects the accuracy of the model to a certain extent, which needs further optimization.

**Table 8.** Prediction error of extreme learning machine prediction model

| Year | Actual value/10,000t | Predicted value/10,000 tons | Relative error (%) | Absolute error/10,000 ton |
| --- | --- | --- | --- | --- |
| 2015 | 25458.8 | 25405.3 | 0.21 | 53.46348 |
| 2016 | 22446.9 | 22229.1 | 0.97 | -217.73493 |
| 2017 | 24327.4 | 24030.6 | 1.22 | -296.79428 |
| 2018 | 22986.3 | 22974.8 | 0.05 | 11.49315 |
| 2019 | 22080.7 | 22071.8 | 0.04 | -8.83228 |
| 2020 | 22237.1 | 22203.7 | 0.15 | 33.35565 |

#### 5.2.3. Prediction based on WOA-BP model

In this paper, the whale algorithm with global search ability is used to optimize the initial weight threshold of BP neural network in order to improve the prediction accuracy of BP neural network. The divided training set and test set are the same as the BP neural network model.

When setting the initial weights and thresholds of the neural network, a set of randomly generated initial values will be selected because there is no relevant setting principle. BP neural network can learn the mapping relationship between input and output automatically, generate initial parameters randomly, and modify the weights and thresholds of the network continuously through error back propagation, but the randomly selected initial weights and thresholds are usually inversely proportional to the convergence speed of neural network training, that is, the larger the value is, the slower the convergence speed is, and then it takes a lot of time. In this case, the final training results are easy to fall into local optimum, and it is difficult to obtain the ideal calculation and prediction results. It can be seen from Table 9 that the relative error of BP neural network after optimization is significantly reduced, which is not more than 1.5%, and the fitting degree of carbon emissions in Guizhou Province is significantly higher than that before optimization. The carbon emission prediction value in 2017 is the closest to the real value, with a relative error of 0.16%, and the prediction results in other years are relatively stable. The accuracy and stability of prediction using WOA-BP neural network have been greatly improved.

**Table 9.** Prediction error of WOA-BP prediction model

| Year | Actual value/10,000t | Predicted value/10,000 tons | Relative error (%) | Absolute error/10,000 ton |
| --- | --- | --- | --- | --- |
| 2015 | 25458.8 | 25260.2 | 0.78 | 198.57864 |
| 2016 | 22446.9 | 22170.8 | 1.23 | -276.09687 |
| 2017 | 24327.4 | 24288.4 | 0.16 | -38.92384 |
| 2018 | 22986.3 | 22834.5 | 0.66 | 151.70958 |
| 2019 | 22080.7 | 21928.3 | 0.69 | -152.35683 |
| 2020 | 22237.1 | 22119.2 | 0.53 | 117.85663 |

**Table 10.** Prediction error of WOA-ELM prediction model

| Year | Actual value/10,000t | Predicted value/10,000 tons | Relative error (%) | Absolute error/10,000 ton |
| --- | --- | --- | --- | --- |
| 2015 | 25458.8 | 25458.8 | 0 | 0 |
| 2016 | 22446.9 | 22303.23984 | 0.64 | 143.66016 |
| 2017 | 24327.4 | 24215.49396 | 0.46 | 111.90604 |
| 2018 | 22986.3 | 22979.40411 | 0.03 | 6.89589 |
| 2019 | 22080.7 | 22078.49193 | 0.01 | 2.20807 |
| 2020 | 22237.1 | 22225.98145 | 0.05 | 11.11855 |

**Table 11.** Comparison of prediction accuracy of different models

| Model category | BP neural network model | ELM model | WOA-BP model | WOA-ELM model |
| --- | --- | --- | --- | --- |
| MAE | 575.93 | 234.46 | 349.65 | 101.13 |
| MAPE | 1.08% | 0.44% | 0.63% | 0.21% |
| RMSE | 761.6 | 338.65 | 387.16 | 166.16 |

**Table 12.**
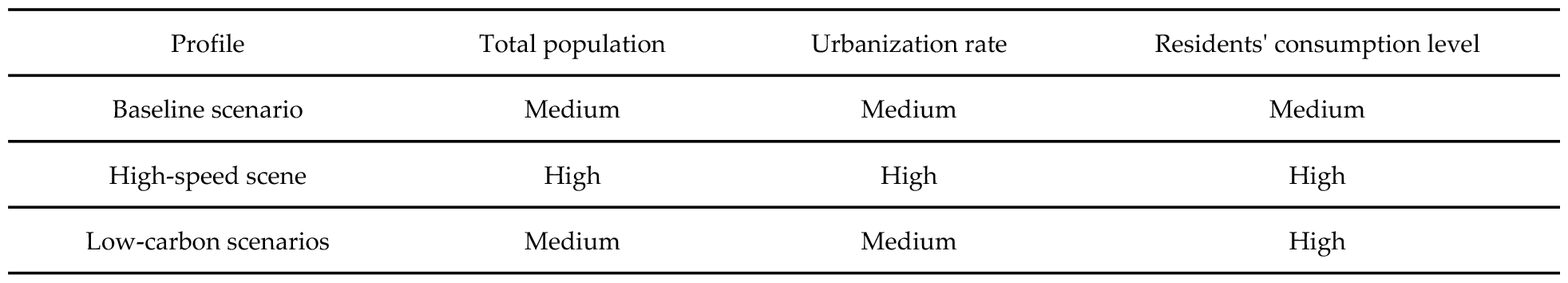
Profile Setting Description

| Profile | Total population | Urbanization rate | Residents' consumption level |
| --- | --- | --- | --- |
| Baseline scenario | Medium | Medium | Medium |
| High-speed scene | High | High | High |
| Low-carbon scenarios | Medium | Medium | High |

**Table 13.** Parameter Settings for Different Scenarios

| Influencing factors | Profile | Rate of change/% |
| --- | --- | --- |
| Total population | Baseline scenario | 0.70 |
|  | High-speed scene | 1.00 |
|  | Low-carbon scenarios | 0.50 |
| Urbanization rate | Baseline scenario | 1.25 |
|  | High-speed scene | 1.50 |
|  | Low-carbon scenarios | 1.00 |
| Residents' consumption level | Baseline scenario | 8.00 |
|  | High-speed scene | 9.00 |
|  | Low-carbon scenarios | 7.50 |
| GDP per capita | Baseline scenario | 6.50 |
|  | High-speed scene | 7.00 |
|  | Low-carbon scenarios | 6.00 |
| Total energy consumption | Baseline scenario | -1.50 |
|  | High-speed scene | -2.00 |
|  | Low-carbon scenarios | -2.50 |

#### 5.2.4. Prediction based on WOA-ELM model

The training samples selected in this section are the same as the extreme learning machine model setting. The input variable and output variable data from 2000 to 2014 are used as the training set, and the prediction year is from 2015 to 2020. The WOA-ELM model is established.

The results show that after multiple training, the ELM model has a better fitting effect on the carbon emissions of Guizhou Province from 2015 to 2020 after WOA optimization, the error between the predicted value and the real value is between 0 and 0.05%, the relative error and absolute error between the two are relatively small except for a few, and the prediction accuracy is significantly higher than that of the ELM model. The effectiveness of the WOA-optimized ELM scheme is verified.

### 5.3. Comparison of prediction results

In this paper, four prediction models were established to test the carbon emission data of Guizhou Province. In order to facilitate the evaluation of the prediction performance of the four models, three indicators of mean absolute error, mean absolute percentage error and root mean square error were used for comparative analysis. The mean absolute error can represent the actual prediction error. The root mean square error represents the deviation between the observed value and the true value.

It can be seen from the above table that the BP neural network is highly accurate in predicting the carbon emission data of Guizhou Province by using the extreme learning machine model. Compared with the ELM model, the BP neural network model is less robust and random, and its convergence speed is slightly lower than the ELM model. Among the four prediction models, the prediction model based on WOA-ELM has the highest prediction accuracy. The prediction model based on extreme learning machine takes the second place. The prediction effect of BP neural network model is the worst. From the above comparison test, it is easy to conclude that the prediction effect of WOA-ELM model is relatively better, and the prediction accuracy can reach the expected level, which can be used to predict the peak carbon emissions of Guizhou Province in the following text.

## 6. Prediction of Carbon Peak in Guizhou Province

### 6.1. Construction of carbon emission scenarios

#### (1) Scenario settings

Scenario analysis refers to the quantitative analysis of the past and present actual situation, which integrates the factors affecting the future and makes qualitative assumptions, so as to infer the possible future situations. It is not the purpose of scenario construction to accurately predict the possibility of the future, but its greatest practical value is to analyze the problem more comprehensively through different ideas. In the process of using scenario analysis, there are two premises, one is to ensure that the impact factors can be quantified, and the other is that the future indicators can be predicted.

This chapter uses the scenario analysis method to set the impact factors of carbon emissions under different development scenarios, so as to predict the level of carbon emissions in Guizhou Province from 2020 to 2040 under various scenarios. The research ideas are as follows: Firstly, three scenarios are set, namely, the baseline scenario, the high-speed scenario and the low-carbon scenario, corresponding to the indicators of medium growth and high growth with positive regression coefficients. Then, according to the strategic policy interpretation of economic development and energy development in Guizhou Province, the current situation of economic and social development and the development trend of energy structure in Guizhou Province are clarified, and the parameters of total population, urbanization rate, residents’consumption level, total energy consumption and per capita GDP in Guizhou Province in the future under different development scenarios are set in combination with relevant policies and energy target requirements. Finally, the evolution trend of carbon emissions in Guizhou Province in the future is predicted.

Benchmark scenario: The benchmark scenario is the continuation of the existing economic development and energy development in Guizhou Province. According to the current economic development model, each factor is set according to the most likely situation. As a large industrial province, Guizhou Province has a very complete industrial infrastructure construction, and its future will also be in the development mode of industrial development for a long time. But at the same time, its economic structure and industrial structure will continue to follow the call of the state for transformation and upgrading. For the energy consumption structure, Guizhou Province, which is dominated by industrial development, will still be dominated by fossil energy consumption, and with the development of new energy technologies becoming more mature, the proportion of fossil energy consumption will continue to decline.

High-speed scenario mode: In the high-speed scenario mode, the five variables of total population, urbanization rate, per capita GDP, household consumption level and total energy consumption all maintain rapid development and change. With the rapid growth of population, the acceleration of urbanization, the rapid and vigorous development of economy and society, the rapid development of new industries and the dominant position of information industry, the use of new energy will be more widely applied to various industries, and the efficiency of energy utilization will be significantly improved.

Low-carbon scenario: the four variables of total population, urbanization rate, per capita GDP and total energy consumption develop at a lower rate than the baseline scenario.

#### (2) Scenario parameter settings

1. Population setting: With the economic and social development, the total population size will continue to expand in the short term, and in the long run, the population growth rate will decline. The analysis of the change trend of the total population of Guizhou Province shows that the population of Guizhou Province is gradually decreasing from 2010 to 2020, and the natural growth rate of the population is negative. By 2020, the population of Guizhou Province is 38.57 million, while the Population Development Plan of Guizhou Province proposes that the permanent population will reach 50 million by 2030, that is, the average annual growth rate is 0.44%. According to the population development plan of Guizhou Province and the population growth in recent years, this paper sets the annual change rate at 0. 7% in the baseline mode, 1% in the high-speed mode, and 0. 5% in the low-carbon mode.
2. Setting of urbanization rate: With the continuous advancement of urbanization in Guizhou Province, the urbanization rate has reached 50.26% in 2018, which is 2.58 times of that in 1995. The average growth rate in the past five years is 1.12%, and the average growth rate in the past ten years is 1.33%. By observing the changing trend of urbanization in the economies of various countries in the world, the urbanization level of Britain and the United States is in a relatively leading position in the process of global urbanization construction, reaching about 80%, while that of other developed countries is about 70%. Compared with the general level of urbanization construction in China, the urbanization process in Guizhou Province is relatively fast. Combined with the experience of developed countries, this paper sets the annual change rate at 1% under the benchmark mode, 1.25% under the high-speed mode, and 1% under the high-speed mode. In the low-carbon mode, the annual rate of change is 0.7%.
3. Setting of per capita GDP: The per capita GDP of Guizhou Province will continue to grow from 2000 to 2020, with the per capita GDP reaching $330/person in 2000 and $7000/person in 2020. In recent years, the development of infrastructure construction has led to the rise of economic development level in Guizhou Province, and the growth rate of per capita GDP has risen rapidly. In 2016, the per capita GDP of Guizhou Province increased by a large margin. According to the 13th Five-Year Plan for National Economic and Social Development of Guizhou Province, the average annual growth rate of regional GDP has reached 6.6%, and the space for the decline of per capita GDP growth rate will gradually shrink after the 13th Five-Year Plan. Therefore, this paper sets the annual change rate at 6.5% under the benchmark mode, 7% under the high-speed mode, and 7% under the high-speed mode. In the low-carbon mode, the annual rate of change is 6%.
4. Residents’ consumption level: From 2000 to 2020, the average annual growth rate of residents’ consumption level in Guizhou Province is 8.0%. According to the “13th Five-Year Plan” of Guizhou Province, it is clearly proposed to release residents’ consumption potential, create consumption demand and further enhance residents’ consumption capacity. Therefore, in the baseline mode, the annual change rate is 8%; in the high-speed mode, the annual change rate is 9%;
5. Total energy consumption: From 2000 to 2020, the total energy consumption of Guizhou Province basically kept a slight increase. From 2002 to 2012, the total energy consumption showed a rapid upward trend. After 2012, the total energy consumption level declined, from 23526 tons of standard coal/10,000 yuan in 2012 to 22321 tons of standard coal/10,000 yuan in 2018. According to the requirements of energy saving and emission reduction planning in the 13th Five-Year Plan of Guizhou Province, this paper sets the annual change rate of-1. 5% in the baseline mode, -2% in the high-speed mode, and -2. 5% in the low-carbon mode.

### 6.2. Prediction and Analysis of Carbon Peak in Guizhou Province

The fitted WOA-ELM is used to predict the carbon emissions of Guizhou Province from 2020 to 2040 under the three scenarios. The predicted results are shown in Table 14.

| Profile | Peak carbon emissions<br>(10,000 tons) | Peak year for carbon emissions<br>(Year) |
| --- | --- | --- |
| Baseline scenario | 26243.61 | 2038 |
| High-speed scene | 30251.37 | 2036 |
| Low-carbon scenarios | 21294.98 | 2033 |

It can be seen from Table 14 that due to the different change ranges of carbon emissions affected by population, urbanization rate, resident consumption level, per capita GDP and total energy consumption, the occurrence time and peak value of carbon peak in Guizhou Province will change due to different parameter settings, and the total carbon emissions will also change accordingly. In the baseline scenario, it is estimated that the peak carbon emissions of Guizhou Province will reach 260 million tons in 2038; in the high-speed scenario, the peak carbon emissions of Guizhou Province will reach 300 million tons in 2036; Under the low-carbon scenario, the carbon peak in Guizhou Province will reach 210 million tons in 2033.By comparing the time and size of the peak carbon emissions in the baseline scenario and the high-speed scenario, it is found that in the baseline scenario, the peak carbon emissions in Guizhou Province will not be achieved in 2030 as scheduled, and will most likely be delayed to 2038. In the high-speed scenario, the peak carbon emissions in Guizhou Province will occur two years earlier than in the baseline scenario. By comparing the time and size of the peak carbon emissions under the baseline scenario and the low-carbon scenario, it is found that the peak year under the low-carbon scenario is earlier than that under the baseline scenario. Although it is three years later than the peak target of China in 2030, the development status of Guizhou Province is relatively backward compared with that of developed cities such as Beijing and Shanghai, so it is acceptable. Its peak volume is 40 million tons lower than the baseline scenario. Comparing the peak time and size of carbon emissions predicted by the low carbon scenario and the high speed scenario, we can see that the peak time of carbon emissions in the low carbon scenario is three years earlier than that in the high speed scenario, and the peak volume is reduced by 50 million tons.It can be seen from the above prediction results that Guizhou Province can not achieve the ambitious goal of carbon peak in 2030 in the baseline scenario and the high-speed development scenario. In contrast, the carbon peak time in the low-carbon scenario is earlier and the peak value is lower.

## 7. Prediction of Carbon Peak in Guizhou Province

Exploring the main factors affecting carbon emissions in Guizhou Province will play an important role in promoting China’s carbon peak and carbon neutralization. In addition, the accuracy of predicting carbon emissions is of great significance for the government to formulate relevant policies and innovate energy saving and emission reduction science and technology. In this paper, firstly, the characteristic subset affecting carbon emissions is constructed by referring to the existing literature and combining with the actual situation of Guizhou Province; secondly, the appropriate input variables are selected based on the grey correlation analysis method; thirdly, the BP neural network and ELM model are established, and the WOA algorithm is used to optimize the BP neural network and ELM model, and the performance of the prediction model is compared and analyzed through simulation. Finally, three scenarios are set to predict the carbon emissions of Guizhou Province from 2020 to 2040. Through the analysis, the following conclusions are drawn:

The results show that the total carbon emissions of Guizhou Province in 2020 will be. 1 million tons 22237, which is about twice the total carbon emissions of Guizhou Province in 2000. With the development of social economy, the growth rate of total carbon emissions in Guizhou Province will gradually decrease, and the overall trend is “S” curve. Secondly, combined with the actual situation of Guizhou Province and previous studies, 12 influencing factors were selected.According to the degree of correlation, the population and total energy consumption have a greater impact on carbon emissions in Guizhou Province.At the same time, the total population, urbanization rate, residents’ consumption level, per capita GDP and total energy consumption were selected as the input variables of the prediction model.

The BP neural network, ELM model, WOA-BP and WOA-ELM models were established to predict the carbon emissions in Guizhou Province. Comparing the mean absolute error, mean absolute percentage error and root mean square error of BP neural network, ELM, WOA-BP and WOA-ELM prediction model, it is found that the accuracy of WOA-ELM model is higher, its MAE is 101. The prediction accuracy of the model based on extreme learning machine is the second, MAE is 224. 46, MAPE is 0. 43%, RMSE is 328. 62, and the prediction effect of BP neural network model is the worst. Three different scenarios are constructed: baseline mode, high-speed mode and low-carbon mode. Using the fitted model and inputting the set scenario parameters into the fitted model, the carbon emissions of Guizhou Province in the next 20 years are carried out. The results show that under the baseline model, the carbon peak of Guizhou Province will appear in 2038, and the peak value will be 0.61 million tons 26243. Under the high-speed scenario, the peak time of carbon in Guizhou Province appears in 2036, and the peak value is 0.27 million tons 30251. Under the low-carbon scenario, the peak time of carbon in Guizhou Province is 2033, and the peak value is 9800 tons 21294. In the baseline model, Guizhou Province cannot achieve the peak target of China in 2030, and the low-carbon scenario is the closest to the carbon peak target of the three scenarios, so it is necessary to intervene in the external policies of Guizhou Province.

According to the results of the above grey correlation analysis, it is found that the population has an important impact on carbon emissions in Guizhou Province, which must be paid attention to in the work of energy conservation and emission reduction in Guizhou Province. Due to the increasing demand for energy in human daily life and production activities, it will promote the increase of carbon emissions and have a significant impact. Controlling the population of Guizhou Province and encouraging citizens to choose green travel and energy-saving and environmentally friendly life will have a far-reaching impact on changing the current situation of carbon emissions in Guizhou Province. Therefore, the government should encourage people to save electricity, do a good job in the disposal of waste household appliances, and increase investment in research and development of energy-saving household appliances. Enrich urban public transport, promote the construction of public transport facilities, and open more convenient energy vehicle development.

In the analysis of the differences in carbon emissions caused by the three development modes in Guizhou Province under the scenario analysis method, it is found that the low-carbon mode is the earliest to reach the highest level of carbon emissions, followed by the high-speed mode, and the benchmark mode is the latest, and it can be found that in the low-carbon development mode, when carbon dioxide emissions reach the peak, the value is the smallest in the three modes. From the overall consideration, population factors, economic development factors and energy consumption factors influence each other, in order to reach the ideal time of carbon peak in Guizhou Province, we should not only ensure the normal economic growth of Guizhou Province, but also take some measures to control the increase of urbanization rate, reduce energy consumption and optimize energy structure. In the energy consumption structure of Guizhou Province, coal consumption accounts for a high proportion, and coal combustion will increase carbon dioxide emissions. Therefore, Guizhou Province should reduce the consumption of coal energy, increase the utilization and conversion rate of coal, increase the investment in research and development of new energy, broaden the scope of popularization of new energy, improve the construction of related supporting facilities, and put the full use of new energy on the agenda. Focus on the development of water conservancy and hydropower projects and photovoltaic projects, and increase the proportion of clean energy such as hydropower in the use of electricity.

## Author Contributions

Conceptualization, W.Y.; methodology, L.D.; software, Y.SQ.; formal analysis, W.Y.; investigation,N.Y.; data curation,WR.R.; writing—original draft preparation, W.Y.; writing—review and editing, N.Y. and L.D.; visualization,Y.SQ.; supervision, WR.R. All authors have read and agreed to the published version of the manuscript.

## Institutional Review Board Statement

Not applicable.

## Informed Consent Statement

Not applicable.

## Data Availability Statement

Not applicable.

